# Urine Recirculation During Normothermic Kidney Preservation Improves Energy Balance Involving the Urea and TCA Cycles

**DOI:** 10.64898/2026.01.12.698950

**Authors:** Annemarie Weissenbacher, Zhanru Yu, Honglei Huang, Maria Letizia Lo Faro, Bohan Yu, James P Hunter, Rutger J Ploeg, Constantin C Coussios, Peter J Friend, Benedikt M Kessler

## Abstract

**Background:** Deceased donor kidneys experience cellular stress before undergoing transplantation. To alleviate this, preservation techniques were developed including normothermic machine perfusion (NMP).

**Methods:** Here, we performed kidney NMP on discarded human kidneys for up to 24 hours. Volume management was regulated either by urine recirculation (UR) or urine replacement (NUR) with Ringer’s lactate. Notably, UR led to longer perfusion times compared to NUR. To investigate kidney NMP metabolic traits with or without UR over time, we performed longitudinal metabolomics analyses of perfusates of eight NMP kidneys by 2D-gas chromatography mass spectrometry (GCxGC-MS).

**Results:** Over 600 metabolic features were profiled, from which 74 were identified and 54 consistently quantified across 26 perfusate samples. Most notably, elevated levels of disaccharides (different isomers), hydroxy-purines, urea, glutamate and amino acids associated with the perfusion factor UR. Moreover, donor estimated glomerular filtration rate (eGFR) correlated significantly with the accumulation of lactate and gluconate. Most strikingly, lactate levels seemed to be more balanced in UR NMP perfusate, which otherwise accumulated rapidly within the first six hours.

**Conclusions:** Kidney preservation by NMP was previously limited to hours. UR-NMP affected kidney energy homeostasis, carbohydrate & purine metabolism and the Urea and TCA cycles. These insights add value to explain how urine-driven adaptations contribute to prolonged kidney function under NMP.

**Research in context:** *Evidence before this study:* Deceased donors provide kidneys that experience cellular stress during retrieval and during transplantation. To attenuate tissue damage, preservation techniques were optimised to offer the most optimal environment for kidney organs retrieved from donation after circulatory death and brain death patients. Normothermic machine perfusion (NMP) of donor kidneys has been considered feasible, safe, offers viability assessment and contributes to favourable outcomes. However, there has been a limit in the length of time that this preservation method could be applied to kidney organs, thereby potentially restricting functional recovery of the kidney before organ transplantation.

*Added value of this study:* Before urine recirculation (UR) was introduced, NMP time was limited to a few hours. Remarkably, NMP with UR led to longer perfusion times and more stable kidney organ function as compared to no urine recirculation (NUR). In order to find out why, we compared urine recirculation with Ringer’s lactate solution for volume management during NMP on discarded human kidneys for up to 24 hours. Kidney NMP metabolic traits with or without UR over time were measured. More than 600 metabolic features were profiled, Most strikingly, lactate levels seemed to be more balanced in UR NMP perfusate of the course of 24 hours, which otherwise accumulated rapidly within the first six hours. Taken together, UR-NMP affected kidney energy metabolic pathways and rendering these more balanced. Ultimately, these urine-driven adaptations contribute to prolonged kidney function under NMP.

*Implications of all the available evidence:* NMP as a procedure to preserve kidney organs including urine recirculation is now becoming standard in many transplantation units around the world. Our study provided a molecular snapshot of why kidney organs preserved in this way are maintained longer with a functionally active metabolism. This provides the basis for additional improvements leading to better kidney organ preservation, ultimately resulting in benefits for kidney transplant recipients.

## Introduction

Kidney transplantation is the preferred treatment option for most of patients suffering from end stage renal disease (ESRD), which is a major global health burden. By 2030, an estimate of 5.4 million people will receive renal replacement therapy (RRT), and it has been projected that seven million adults will have died due to lack of access to RRT ^1^. Access to kidney transplantation is limited though, as there is evident organ scarcity and a major disparity between transplantable kidneys and patients undergoing RRT on the waiting lists. Enlarging the donor pool may be achieved by recovering and transplanting more marginal organs from extended criteria (ECD) and donors after circulatory death (DCD), the fastest growing source of organs ^2^.

Organ preservation has been crucial for organ transplantation. During the last two decades, the focus has been on the optimisation of preservation techniques to offer the most optimal environment for ECD and DCD organs. Since the landmark study has described the favourable outcome in terms of lower rates of delayed graft function (DGF) after kidney transplantation following hypothermic machine preservation (HMP) compared to static cold storage (SCS), HMP has been implemented in the clinical routine in many kidney transplant units ^3–5^.

Implementing kidney preservation technologies like hypothermic machine perfusion (HMP) ^3^ and moving kidney viability assessment to a clinically applicable level by normothermic machine perfusion (NMP) of deceased donor kidneys ^6–8^ have been relevant measures contributing to increasing the supply of transplantable kidneys. The term ‘transplantable kidney’ is a crucial one, and the main reason why the community wants to apply kidney NMP, where affordable, to fill the void between currently available organs and transplantable recipients on the institutional waiting list.

In the clinical trial setting, kidney NMP has been applied for one to two hours before such organs have been either declined or transplanted ^6,9^. Investment in dynamic kidney NMP offers the ability to create a near-physiological condition for the organ. Under these circumstances, the kidney starts to function, viability assessment becomes possible and repair mechanisms are initiated ^7,10^.

Several ways to perfuse a kidney normothermically and to assess viability have been described ^11^. (Urine recirculation (UR), in comparison to urine replacement (NUR), leads to significantly longer NMP times and also to an improved homeostatic state of the perfused organ ^12,13^. The perfusate, however, which is an absolutely crucial parameter in kidney NMP, has not been investigated yet in terms of differences in its metabolome related to volume management of the excreted urine. Perfusate metabolomics analyses in kidney HMP show the evolution of metabolites and their increased imbalance during the preservation period ^14,15^. Furthermore, metabolic differences between immediately functioning kidneys after HMP and kidneys with delayed graft function after HMP have been described ^16,17^. Metabolomics technology has also been applied to study small molecules affected by the detrimental effects of ischemia-reperfusion injury ^18^. However, in clinical transplantation or in pre-clinical NMP trials evaluating human kidneys, the metabolic process has not yet been analysed and described extensively.

The primary aim of our analysis was to investigate the metabolome and its potential change over time in the kidney perfusate NMP conditions. Moreover, it was of interest to detect any differences between UR and NUR, as well as correlations with kidney donor demographics and perfusion characteristics that may contribute to prolonged perfusion with UR.

## Methods

Discarded human kidneys were normothermically perfused for up to 24 hours on a prototype device for normothermic machine preservation (NMP) of the kidney as described previously ^12,19^. A blood-based perfusate was used using one unit of packed red blood cells (RBCs) of the same blood group as the kidney that was resuspended in 250 mL of 5% human albumin solution as reported in detail ^12,19^. RBCs used were quality tested by the blood bank. Supplements were added prior to the initiation of kidney NMP; 750 mg cefuroxime, 10 ml calcium gluconate 10%, 80 mg enoxaparin, 5-15 ml of sodium bicarbonate 8.4% to reach a perfusate pH of 7.3 under physiological pCO_2_ conditions of 4-6 kPa ^12,19^. Epoprostenol sodium was infused at a constant rate of 4 μg/h during kidney NMP. Bolus injections of lipid-free parenteral nutrition solution (0.24 g/ml glucose) or glucose 5% (0.05 g/mL glucose) were administered in 2.5-5 mL increments once blood glucose levels dropped below 4 mmol/L. For perfusate volume control, recirculation of the urine (UR – subjects 4, 7, 21, 28 and 35) or replacement of the urine (no urine recirculation, NUR – subjects 29, 31 and 32) with Ringer’s lactate was applied ^12,13,19^.

### Ethics statement

Kidney retrieval and NMP procedures were approved by the national ethics review committee of the United Kingdom (REC reference 12/EE/0273 IRAS project ID 106793).

#### Extraction of metabolites from perfusate samples

Two hundred μL of perfusate collected at the beginning of perfusion, after six hours (NUR and UR), after twelve and 24 hours of perfusion (kidneys with UR) were used for metabolites extraction. The perfusate sampling protocol and the metabolomics workflow are illustrated in **Figure 1**. Two hundred μL methanol were added to the perfusate and mixed for 5 minutes on a shaker at 1500 rpm. Five μL myristic acid-14,14,14,d3 (1mg/mL, Cambridge Isotope Laboratories, United Kingdom) was spiked into perfusate samples as normalisation control. The samples were vortexed for 5 minutes then followed by adding of 1 mL of *tert*-butyl methyl ether (MTBE). The samples were mixed by shaking for 5 minutes then centrifuged for 20 minutes at 13,000 g at 4°C. The top layer organic phase (MTBE) was transferred into a glass vial and dried in a Speed Vac (Eppendorf, United Kingdom). Subsequently, 800 μL of methanol were added into the aqueous remain (bottom layer), the samples were vortexed for 5 minutes and centrifuged for 20 minutes at 13,000 g at 4°C. The supernatant (aqueous phase) was collected and added into the glass vial containing the organic phase to dry under vacuum. The dried samples were kept at -80°C until use ^19^.

**Figure 1:**
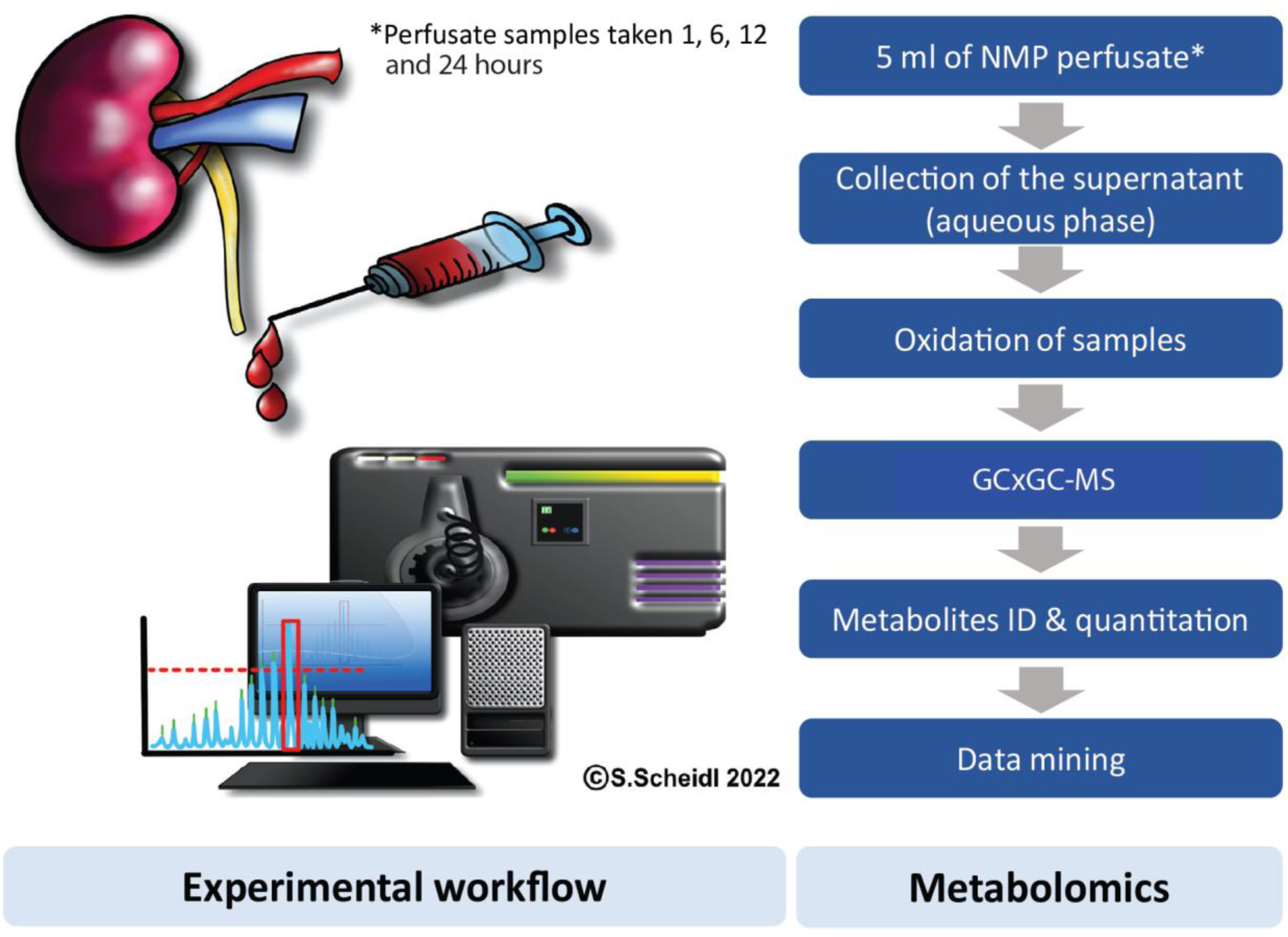
Experimental workflow of kidney NMP perfusate metabolomics analysis. Aliquots of normothermic perfusion (NMP) perfusate (+/- urine recirculation UR versus NUR) were collected after 1, 6, 12 and 24 hours (hrs), respectively. Samples were subjected to organic extraction, chemical derivatization and analysed by two-dimensional gas chromatography mass spectrometry (GCxGC-MS).

Chemical derivatisation was performed essentially as described ^20^. In brief, samples were oxidised in 50 μL of methoxyamine hydrochloride in pyridine (20 μg/µL) and shaken (1200 rpm) for 90 minutes at 30°C. Seventy μL of N-Methyl-N-trimethylsilyltrifluoroacetamide (MSTFA) with 1% chlorotrimethylsilane (TMCS) and 30 µL pyridine were added to the samples, followed by incubation for one hour at 60°C at a shaking speed of 1200 rpm. The samples were cooled down to temperature ambient and injected directly for GCxGC-MS analysis ^19^.

#### Metabolomic analysis using gas chromatography mass spectrometry (GCxGC-MS)

Analysis of perfusate metabolites was performed essentially as described ^20,21^. In brief, two µL of derivatised metabolites were injected into a two-dimensional gas chromatography single quadrupole mass spectrometer (GCxGC-qMS) instrument (GP2010, Shimadzu, United Kingdom) with a split ratio of 1 in 20. The oven temperature was programmed to rise from 80°C to 330°C at a rate of 6°C per minute, and the total run time was 35 minutes. The metabolites were first separated by a non-polar column (30m x 0.25mm inner diameter), followed by a second polar column (6m x 0.25 mm inner diameter) controlled by a modulator that operated at six second cycles as described ^22^. In brief, metabolite effluents separated on the first column were accumulated for 6 seconds, then remobilised, focused and injected into the second column. Ions were generated within GCxGC-MS by electron impact (EI) ionisation and collected at a mass range of 50*m/z* to 800*m/z* using an acquisition rate of 20,000 u/sec and 100 Hz ^19^.

### Metabolomics data analysis

GCMSsolution software (v2.72/4.20 Shimadzu) was utilised to initially process raw GCxGC MS data. A representative sample with a maximal number of peaks present in the data was chosen to act as a reference to ensure comprehensive coverage of metabolites. Chromsquare software (v2.1.6, Shimadzu) was then employed to identify metabolites present in the sample. Metabolites were identified by matching collected fragment ions to the National Institute of Standards and Technology (NIST 11/s, OA_TMS, FA_ME) and YUTDI in-house libraries as references. All other samples were subsequently processed to ensure that no metabolites were missing from the reference sample. The identification confidence score cut-off was set to 80 (100 max.). A panel of collected standards including palmitate, stearate, glucose, maltose, saccharose, lactose and mannose, were used to verify metabolite identifications. Metabolites were identified as chemically derivatised forms, often including several different derivative moieties (methyloxime or trimethylsilyl groups), thereby increasing confidence of identification (**Table 2**). This list of metabolites was used as a key for the development of Python-based scripts to filter the ‘blobs’ identified by Chromsquare in each sample. To identify a blob, its first retention time was required to match the times up to two modulation steps (of 6 s) away from the time in the key (within an error of 0.12 s in each case). Additionally, either the blob was required to have already been identified by Chromsquare or to have a second retention time within 0.2 s of the metabolite list. For each sample, the areas reflecting the same metabolite were then summed. For data curation, the retention times in the metabolite list were manually verified regarding overlapping retention times of several metabolites that fell within the area assigned to a blob by the Chromsquare software.

For metabolite quantitation, mass intensities measured for different chemical derivatives reflecting the same metabolite were combined and analysed using Chrome square software 2.1.6 (Shimadzu, Japan). For the metabolites citrate, succinate and phosphoglycerol, the characteristic fragment ions (m/z values 273, 247, 299, respectively) were used for quantitation within the GCMSSolution software. In these cases, the metabolites were represented by their relevant m/z values and retention times (15.741, 9.942 and 15.041 min., respectively). In-house Python scripts were written to sum the areas of peaks with matching m/z values and retention times. The retention time was required to match the times up to one modulation step (of 6 s) away from the time in the key (within the same error as for the first retention time previously). Each metabolite quantity was normalised to spiked deuterated myristic acid-14,14,14-d3 for comparison as described previously ^19^. Saturated detection levels by GCxGC-MS were observed for several metabolites including lactate and glucose, which was taken into consideration during the MS quantitation data processing step.

### Statistical analysis

For statistical evaluation of the data, excluding metabolomics, unpaired t tests (parametric) and Mann-Whitney tests (nonparametric) were performed using GraphPad Prism 9. Correlation analyses were performed using IBM® SPSS® Statistics Version 28. A p-value of less than 0.05 was considered significant. Hierarchical clustering analysis, heatmap and volcano plots were generated using Perseus software (v1.6.0.2; https://maxquant.net/perseus/). Pearson correlation analysis was used for both metabolites and patient samples, and a student t test was used for the volcano plot analysis, applying Permutation FDR for multiple test correction ^23^. Metabolite and enrichment analyses were performed using MetaboAnalyst v6.0 (https://www.metaboanalyst.ca/). To this end, metabolomic data generated by Chromsquare (see above) were imported into MetaboAnalyst using metabolite identities and associated peak intensities. The normalisation was performed based on the sum. Data was processed by log transformation (base 10) and ‘auto scaling’ for data scaling and visualisation. To replace missing values, we used limit of detection (L.o.D.) 1/5 of the minimal positive value per metabolite as default option offered within the MetaboAnalyst software. Correlation (**Figure S1A**) and box plot analyses (**Figure S1B**) were carried out using Instant Clue software (version 5.2; http://www.instantclue.uni-koeln.de/).

## Results

Eight discarded human kidneys underwent NMP up to 24 hours (**Figure 1**), 5/8 (62.5%) with urine recirculation (UR) and 3/8 (37.5%) with urine replacement using Ringer’s lactate (NUR) (**Table 1**). Seven (87.5%) kidneys were from donors after brain death; the median (min-max) kidney donor age was 72 (61-78) years. Median (IQR) donor eGFR at retrieval was 91 (49.5) mL/min/1.73m^2^. Overall median (IQR) cold ischemia time was 19.9 (4.1) hours; 19.1 (47.9) hours for UR and 21.1 (3.6) hours for NUR kidneys. Overall median (IQR) duration of NMP was 24 (15.6) hours; 24.1 (0.2) hours for UR and 8.2 (3.3) hours for NUR kidneys. Median (IQR) renal arterial flow during NMP was 392 (237.3) mL/min; 413 (370.7) ml/min in kidneys with UR and 371 (386) mL/min in kidneys with NUR as volume control. Median (IQR) urine excretion per hour during NMP was 52.3 (66.7) ml; 41 (46.1) mL in UR and 84.4 (28) mL in NUR kidneys, respectively. Five of eight kidneys received glucose 5% only as nutrition during NMP, 3/8 received lipid-free parenteral nutrition. Median (min-max) volume of nutrition administered during NMP in UR kidneys was 45 (18-91) mL and 11 (10-30) ml in NUR kidneys, accounting for 4.6 (3.2-10.8) gram glucose in UR and 0.6 (0.5-1.5) gram glucose in NUR kidneys during the entire preservation period.

**Table 1:**
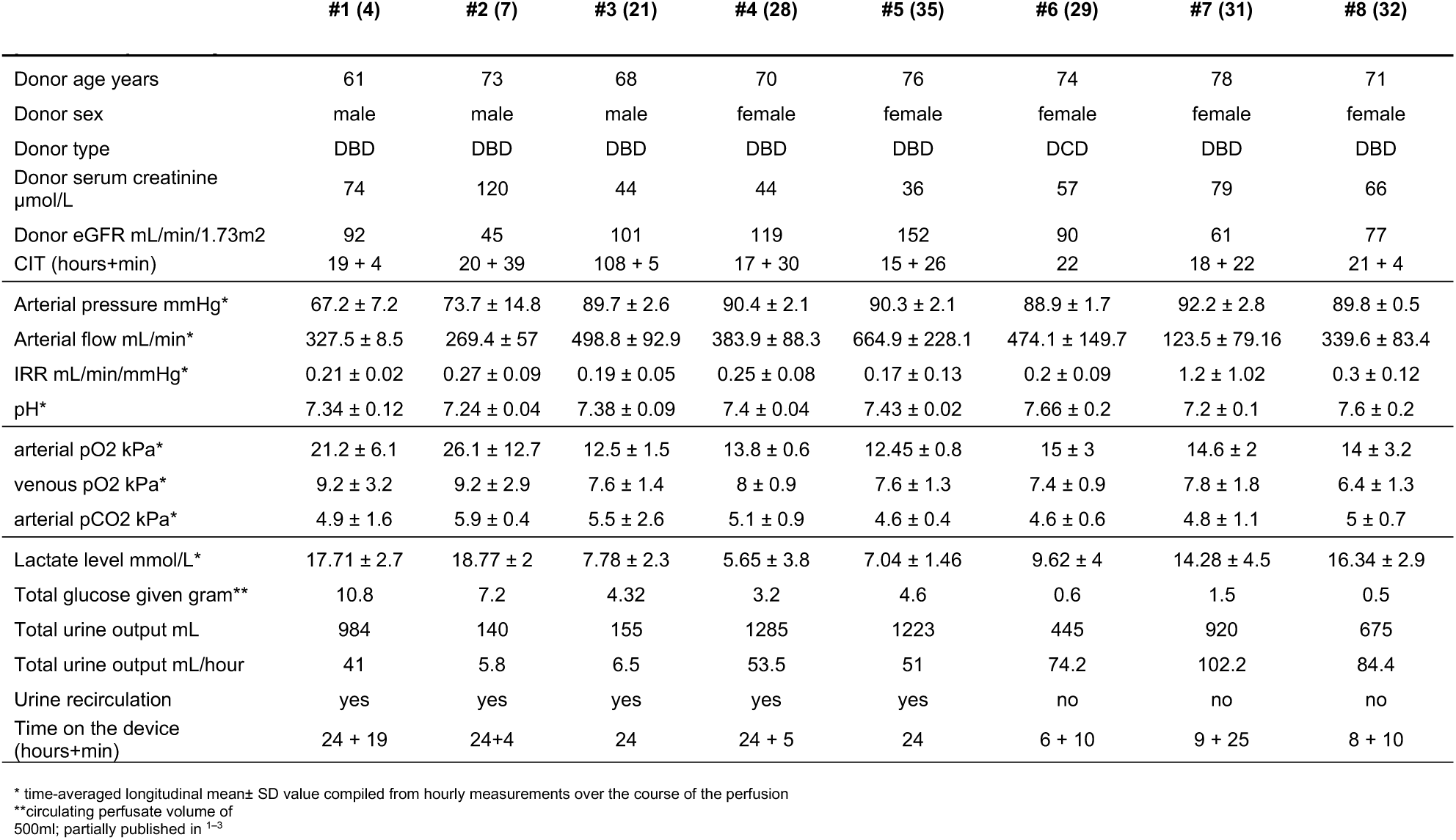
Demographics, hemodynamic and metabolic function parameters per kidney used in this study.

### Kidney perfusate metabolite signatures reflect glycolysis, TCA, urea cycle, β-oxidation, amino acid and purine metabolic pathways

Circa 600 mass features were detected by mass spectrometry, from which 74 metabolites were identified with >80% confidence, most of which with multiple chemically modified derivatives (methyloxime or trimethylsilyl derivatisation). Fifty-four metabolites were confirmed in identity and consistently quantified across 26 perfusate samples, 20 from UR and 6 from NUR kidney perfusions, respectively. The perfusate metabolome comprised mainly glycolysis / citric acid (TCA) metabolites, amino acids and sugars besides organic acids, fatty acids, purines or carbamides (**Figure 2A**) ^19^. All detected and quantified metabolites per group are listed in **Table 2**. For each metabolite, four technical replicates were generated per time point and per sample. These replicates were averaged for quantitative analysis. We first compared metabolic profiles between perfusates with (UR) and without urine recirculation (NUR). Second, metabolite abundances were compared between the different timepoints in both groups, with and without UR. In a third step, differences in metabolites between one-hour perfusate samples of both groups and six-hour perfusate samples of perfusions with and without UR were compared ^19^.

**Figure 2:**
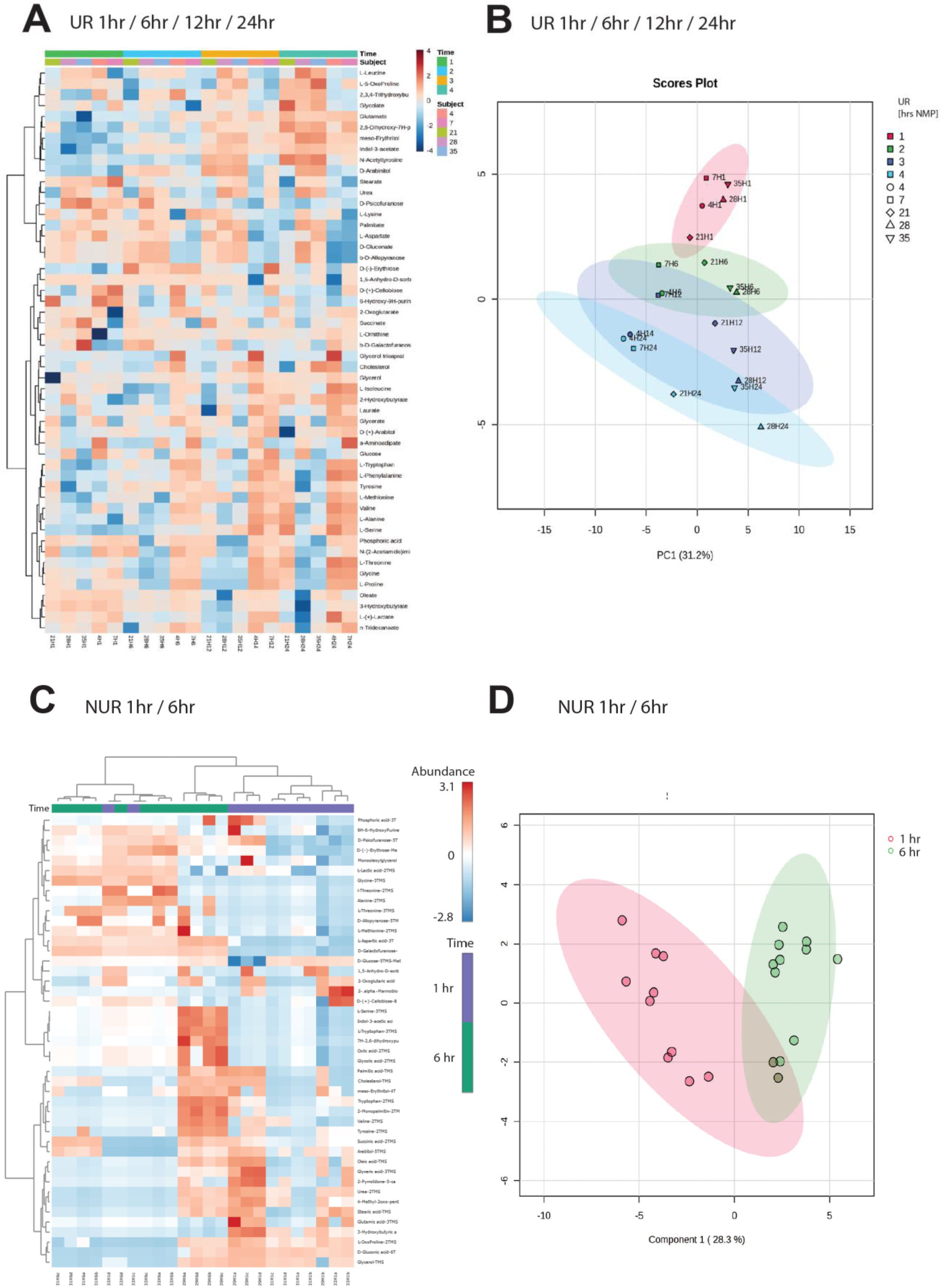
UR alters metabolic profiles and prolongs kidney NMP. (**A**) Hierarchical clustering analysis and heatmap of NMP UR perfusate at 1hr, 6 hrs,12 hrs and 24 hrs. (**B**) Principle component analysis (PCA) of NMP UR perfusate collected at 1 hr, 6 hrs,12 hrs and 24 hrs, respectively. (**C**) Hierarchical clustering analysis and heatmap of NMP NUR at 1 hr (purple bar) and 6 hrs (green bar). Subjects 29, 31 and 32 and four technical replicates per time point (a-d). (**D**) PCA analysis of NMP UR perfusate collected at 1 hr and 6 hr time points including four technical replicates each, respectively.

**Table 2:** Metabolites detected in 26 perfusate samples of normothermic kidney preservation with UR and NUR. Columns represent metabolite groups and native metabolite names.

| Metabolite Group | Metabolite name <sup>[1]</sup> | Metabolite group | Metabolite name <sup>[1]</sup> |
| --- | --- | --- | --- |
| Amino acid | N-acetyltyrosine | Organic acid | 2,3,4-trihydroxybutyrate |
| Amino acid | L-alanine | Organic acid | 2-Hydroxybutyrate |
| Amino acid | L-Aspartate | Organic acid | 2-Oxoglutarate (α-Ketoglutarate) |
| Amino acid | L-Glutamate | Organic acid | 3-Hydroxybutyrate |
| Amino acid | Glycine | Organic acid | 3-Phosphoglycerol |
| Amino acid | L-Leucine | Organic acid | 4-Methyl-oxovalerate |
| Amino acid | L-Isoleucine | Organic acid | α-aminoadipic acid |
| Amino acid | L-Lysine | Organic acid | N-(2-Acetamido)iminodiacetic acid (ADA) |
| Amino acid | L-Methionine | Organic acid | Citrate |
| Amino acid | L-Ornithine | Organic acid | D-Gluconate |
| Amino acid | L-5-OxoProline | Organic acid | Glycerate |
| Amino acid | L-Phenylalanine | Organic acid | Glycolate |
| Amino acid | L-Proline | Organic acid | Indol-3-acetate |
| Amino acid | L-Serine | Organic acid | L-(+)-Lactate |
| Amino acid | L-Threonine | Organic acid | Succinate |
| Amino acid | L-Tyrosine |  |  |
| Amino acid | L-Tryptophan | Inorganic acid | Phosphoric acid |
| Amino acid | L-Valine |  |  |
| | | Sugar | β-D-Allopyranose |
| Fatty acid | Laurate (Dedocanonic acid) | Sugar | 1,5-Anhydro-D-sorbitol |
| Fatty acid | Oleate (oleic acid) | Sugar | D-Arabinol |
| Fatty acid | Palmitate (palmitic acid) | Sugar | D-(+)-Arabitol |
| Fatty acid | Stearate (stearic acid) | Sugar | D-(+)-Cellobiose |
| Fatty acid | n-Tridenoate (n-Tridecanoic acid) | Sugar | D-(-)-Erythrose |
| | | Sugar | β-D-Galactofuranose |
| Purine | 2,6-dihydroxy-7H-purine | Sugar | D-Glucose |
| Purine | 6-hydroxy-9H-purine | Sugar | Glycerol |
|  |  | Sugar | Glycerol |
| Lipid | Cholesterol | Sugar | Meso-Erythritol |
| Lipid | Glycerol tricaprates | Sugar | D-Psicofuranose |
| Carbamide | Urea | External standard | Myristic acid-d3 |

\*Measured in five kidney perfusions with urine recirculation (UR, n=20 samples) and in three kidney perfusions without urine recirculation (NUR, n=6 samples); overall circulating volume was ~500ml perfusate containing ~250ml red blood cells (RBC) and ~250ml 5% albumin)
[1] Metabolites were detected as trimethylsilyl- and/or methyloxime- derivatives by GCxGC-MS. For those metabolites with multiple chemical derivatives detected, MS data was merged (see also Materials & Methods section).

### Correlation of perfusion characteristics and donor factors with metabolites

To investigate a possible relation of donor and perfusate factors (**Table 1**) with the UR/NUR kidney perfusate metabolic profiles (**Figure 2**), correlation analyses were performed with metabolites (**Table S1A**). A major hallmark was that the higher the donor eGFR values of perfused kidneys, the lower the content of lactate and gluconate were noted in perfusates. Duration of CIT prior to NMP-start did not correlate with any of the metabolites in the perfusate; neither in UR nor in NUR kidneys. Duration of NMP, independently of UR and NUR, correlated significantly with L-tyrosine and phosphoglycerol (**Table S1A**). The higher the hourly urine output during NMP, the lower were levels of L-tyrosine and phosphoglycerol in the perfusate of UR and NUR kidneys. Urine recirculation correlated significantly with an elevated presence of disaccharides (D-(+)-cellobiose), gluconate and phosphoglycerol as part of the perfusate metabolome. Lactate showed a significant inverse correlation with median renal arterial flow during NMP. Also, the higher the median (IQR) pCO_2_ levels, 5 (0.53) kPa, were during kidney NMP, independent of UR or NUR, the more lactate levels were found to be elevated in the perfusate (**Table S1A**).

### Altered perfusate metabolite profiles over time of perfusion

Ten metabolites, including three amino acids, two organic acids and two sugars, significantly changed in abundance in perfusates of UR kidneys over time (**Figure 2A,B** and **Table S1B,C**). In particular, for the organic acid lactate, and the amino acid L-alanine, significant changes could be detected during the early phase of perfusion in both, NUR and UR conditions (hour 1 vs hour 6 p=0.004, hour 1 vs hour 12 and hour 24 p<0.0001) (**Table S1B**). For the amino acids L-serine (p=0.001) and glycine (p=0.04), we observed significant changes at later stages of perfusion (up to 24 hours). Similar profiles were noted for the organic acid D-gluconate (p=0.001) and the sugars β-D-allopyranose (p<0.0001) and D-glucose (p=0.002) (**Figure 2A, Table S1B**).

Three metabolites in the perfusates of kidneys with NUR changed significantly between 1 and 6 hours after the start of perfusion (**Figure 2C/D** and **Table S1B**). One organic acid, 3-hydroxybutyrate (p=0.0003), changed significantly. The sugar β-D-galactofuranose (p<0.0001) and the amino acid L-aspartate (p<0.0001) also changed significantly from perfusion start until hour 6 (**Table S1C**). Interestingly, none of the kidney perfusions with NUR reached the later measurements points at hour 12 or hour 24 as observed for the kidneys with UR perfusion, suggesting the existence of substantial metabolic differences between UR and NUR NMP.

### Distinct kidney perfusate metabolite profiles between UR and NUR NMP

To explore metabolic differences due to urine recirculation in detail, we generated a correlation plot and a principal component analysis (PCA), confirming a clear and distinct difference in the global kidney perfusate metabolome of UR and NUR perfusates (**Figure 3A, S1)**. In particular, the perfusate metabolome of UR and NUR kidneys showed distinct differences already after one hour of NMP (**Figure 3A**, B). For instance, the amounts of three sugars, two purines and one carbamide were significantly higher in UR perfusates as compared to levels measured after one hour of kidneys with volume management facilitated by replacing the urine (NUR) with Ringer’s lactate. These include D-(+)-cellobiose (p<0.00001), 2,6-dihydroxy-7H-purine (p<0.00001), 6-hydroxy-9H-purine (p<0.00001) and urea (p=0.00003) (**Figures 3 and 4**). The statistical results for the differences after one hour are displayed in **Table S1B**. Eight metabolites did not change over time at all; neither in the UR perfusate nor in the NUR perfusate cohort (**Table S1 –** also to some extent visible in **Figure 3**).

**Figure 3:**
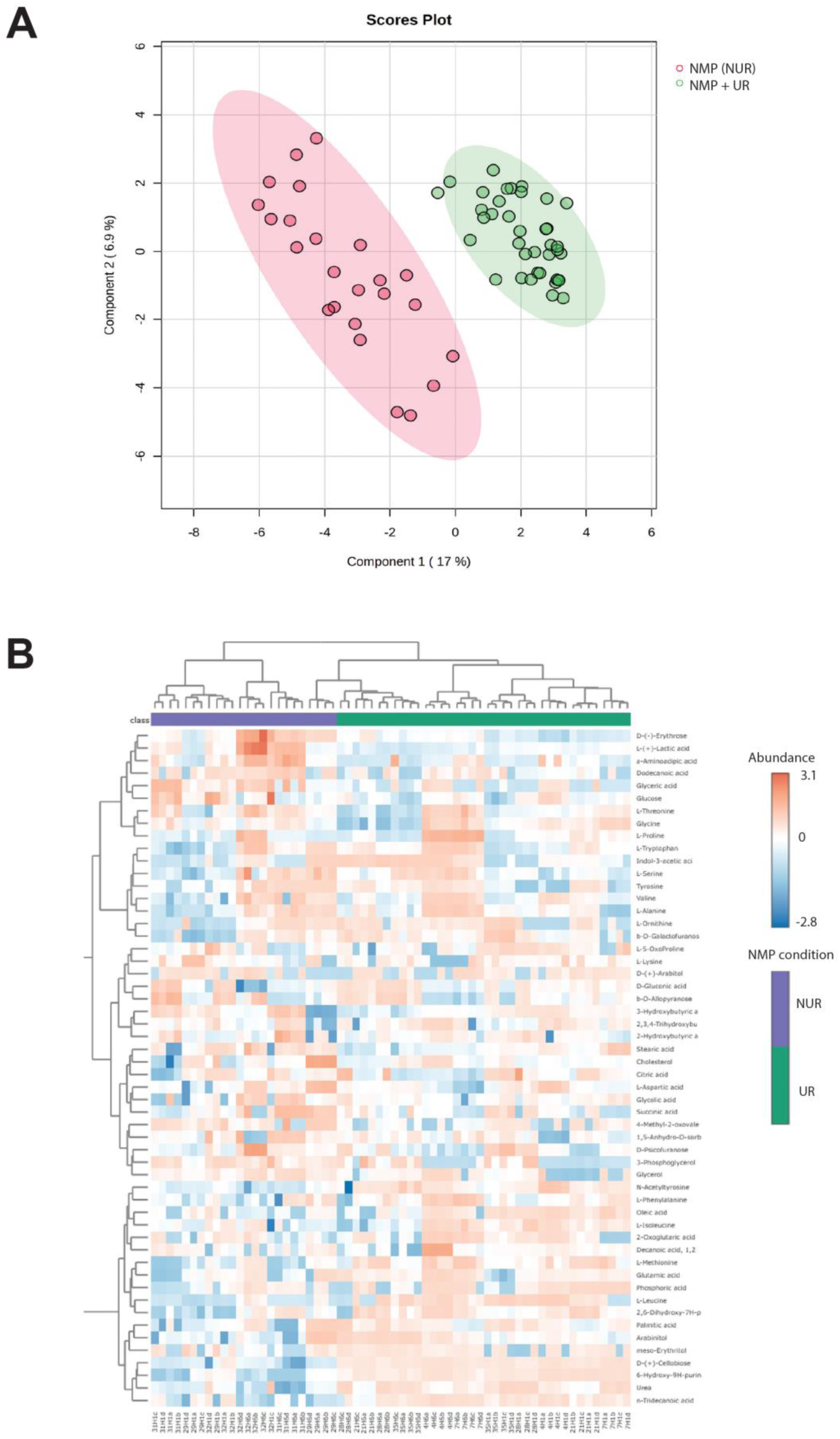
UR modulates kidney NMP metabolic profiles. (**A**) PCA analysis of NMP UR versus NUR perfusate material across all time points including four technical replicates. (**B**) Hierarchical clustering analysis and heatmap of NMP UR versus NUR perfusate across all time points including technical replicates (indicated as a-d).

**Figure 4:**
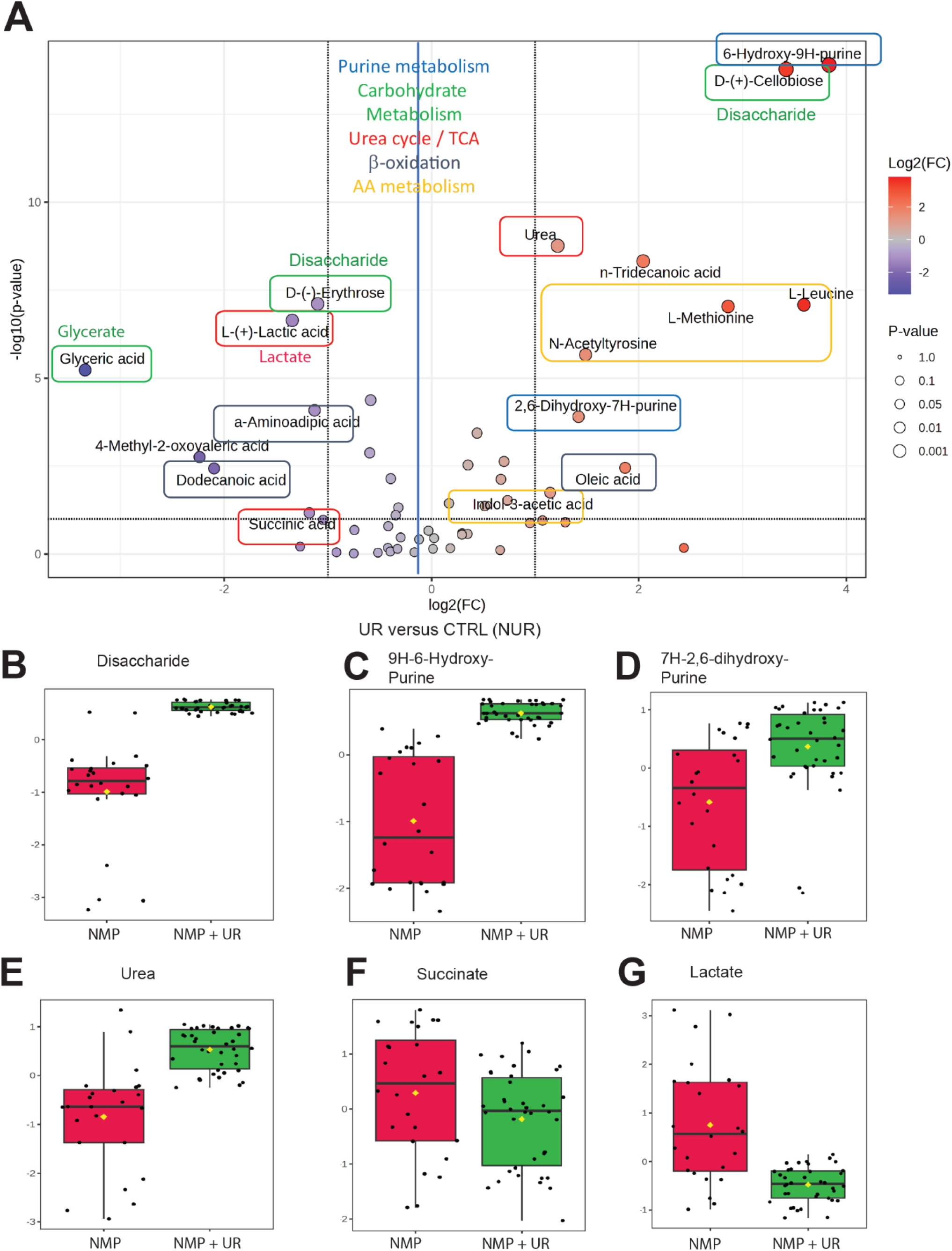
UR results in increased urea cycle, carbohydrate and purine metabolic profiles in kidney NMP perfusate. (**A**) Volcano plot analysis of kidney NMP UR versus NUR perfusate. Violin plot analyses of disaccharide (**B**), 9H-hydroxy purine (**C**), 7H, 2,6-dihydroxy purine (**D**), urea € and glutamic acid (**F**) levels in control (ctrl/NUR - red) versus UR (green) NMP.

After six hours of NMP, there were still remarkable differences between the perfusate metabolome of UR as compared to NUR kidneys. The results for the sugar content of the perfusates were very equivalent to the findings after one hour of NMP. There was one more prevalent amino acid (glutamate, p<0.00001), which could be due to the administration of lipid-free parenteral nutrition solution in the UR kidneys, which contains amino acids. After six hours of NMP, there was not such a striking difference between the presence of purines any longer. The statistical results for the differences after six hours of kidney NMP with either UR or NUR are shown in **Table S1B**.

A volcano plot (student t test) analysis revealed globally distinct kidney perfusate metabolic profiles when perfused normothermically with UR and NUR, assessed at the 1- and 6-hour time points (**Figure 4 A**). In particular, we noted an enrichment of disaccharides in perfusates from kidneys with intact function under UR, clearly different from monomeric glucose by retention time (**Figure S2**). Its MS fragmentation profile (**Figure S3**) revealed the closest match to D-(+)-cellobiose (**Figure S4**), also known as GLCB1-4GLCB or cellose, in kidney perfusates under UR (**Figure 4B**). In addition, we noted accumulation of purines 2,6-dihydroxy-7H-purine and 6-hydroxy-9H-purine (**Figure 4 C,D**), urea (**Figure 4E**) and succinate (**Figure 4F**) in kidney perfusate with UR NMP. Lactate levels were consistently lower in UR versus NUR perfusate (**Figure 4A, G**), a trait that persisted up to 24 hours, although with some kidney specific variation (**Figure 5A**). Interestingly, marked changes in UR perfusate metabolites including glucose, disaccharide (cellobiose) and urea remain remarkably stable during the entire duration of NMP (**Figure 5B-D**). Interestingly, metabolite levels appear to establish themselves at one hour after exposure to UR already (**Figure 5E-H**), and UR promotes a more balanced lactate/glucose ratio (**Figure 5E,F**). Together, we observed considerable metabolic changes between UR and NUR NMP that may suggest possible alterations in carbohydrate, purine metabolic pathways via the Urea - TCA cycle network (**Figure 6**). These changes ultimately contribute to the improvement of the physiological metabolic state of perfused kidney organs.

**Figure 5:**
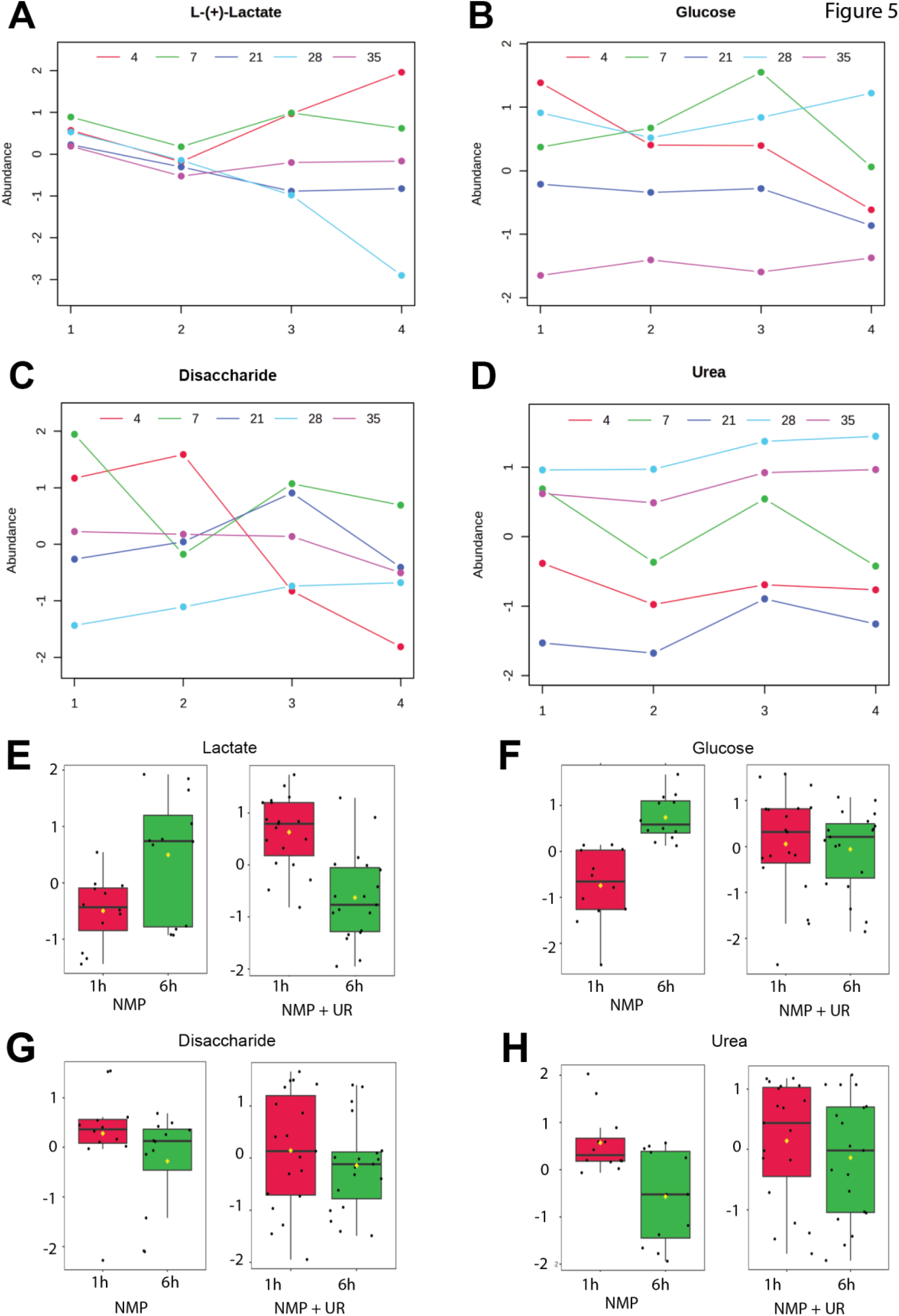
UR NMP perfusate metabolite time course profiles. Abundance levels in UR normothermic perfusate is shown for metabolites lactate (**A**), glucose (**B**), D-cellobiose / disaccharide (**C**) and urea (**D**). Time points 1, 2, 3 and 4 refer to perfusion times of 1, 6, 12 or 14 and 24 hours, respectively. Data for individual kidneys are shown: kidney 4 (red), 7 (green), 21 (dark blue), 28 (light blue) and 35 (purple).

**Figure 6:**
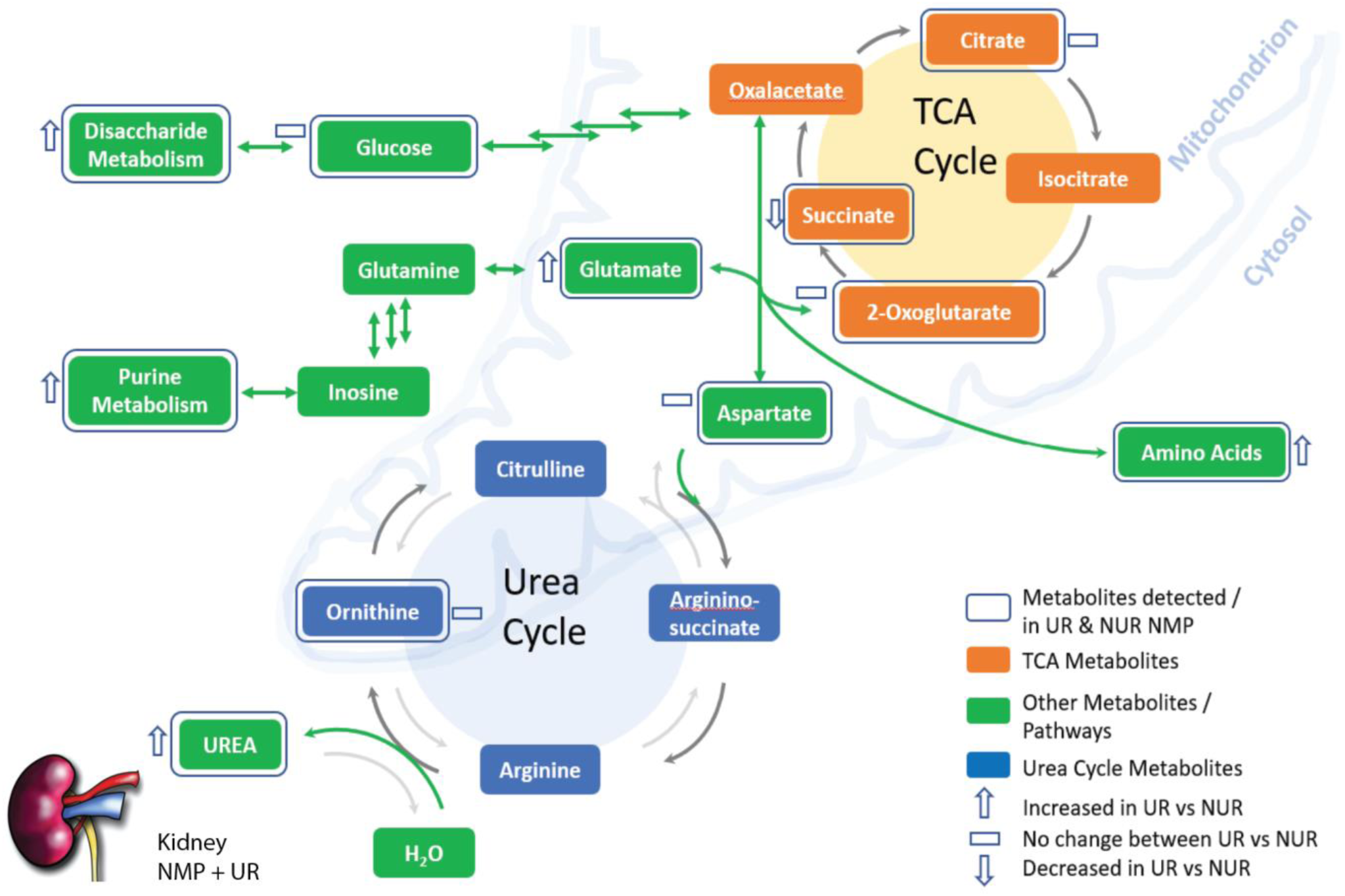
UR NMP may trigger a rewiring of central energy metabolic pathways in perfused kidneys. Enhanced levels of urine (and subsequently urea) in kidney NMP UR may affect the Urea cycle with potential consequences on carbohydrate and purine metabolic profiles. This may occur via direct links between the Urea - TCA cycles and associate metabolic pathways. Molecular details of these interconnections are currently not understood. As a possible key step, glutamate transamination may be leading to oxalacetate to become increasingly available for TCA and carbohydrate metabolisms and may be feeding amino acid and purine anabolism. Ultimately, these metabolic alterations contribute to a more balanced metabolic state of the kidney, resulting in prolonged NMP duration and longer preservation times when urine recirculation (UR) is included.

## Discussion

Ischemic insult and reperfusion injury hampers function of transplanted kidneys. Mitochondrial dysfunction accounts for a major alteration in energy metabolism, mainly due to the disruption of enzyme complexes by reactive oxygen species (ROS), oxygen damage and proteolysis ^21,24^. Evidence for protein breakdown as a hallmark of kidney tissue injury during ischemia reperfusion has been observed previously. For instance, in brain death, cerebral injury contributes to significant cellular stress in donor kidneys adversely impacting the quality of grafts ^25^. Using mass spectrometry and immunoblotting, degradation profiles of cytoskeletal proteins in deceased and living donor kidney biopsies we mapped, α-actinin-4 and talin-1 proteolytic fragments were detected in brain-death donor kidneys caused by calpain activation, subsequently leading to dysregulation of the actin cytoskeleton in human podocytes. In a different study, doxycycline induced changes of the kidney tissue degradome were examined during NMP, yielding a lower level of Neutrophil Gelatinase-Associated Lipocalin (NGAL) in kidneys with less marked tissue degradation protein profiles ^26^. A marked improvement of kidney function and recovery during NMP was achieved by the introduction of urine recirculation with an apparent stabilisation of metabolomic homeostasis and improved electrolyte balance, leading to substantially longer perfusion times ^12,13,19,27,28^. Since proteomics profiles in normothermically perfused kidneys with UR indicated a marked stabilisation of energy metabolism, and to better understand the molecular basis underlying improved organ functionality with UR, we conducted a metabolomics experiment. Perfusate material was subjected to organic extraction, chemical derivatisation and analysis by 2-dimensional gas chromatography single quadrupole mass spectrometry (GCxGC-qMS) ^20,21^.

The results of the metabolomics analyses of these eight discarded human kidneys undergoing NMP for up to 24 hours, applying either UR or NUR, to manage the volume of excreted urine, revealed a distinct pattern of metabolites showing a clear difference between urine recirculation or replacement with Ringer’s lactate. In particular, we noted an enrichment of disaccharides in perfusates from kidneys with intact function under UR. The exact nature of the disaccharide was explored further by comparing MS profiles to standards including glucose (**Figure S4A**), lactose (**Figure S4B**), maltose (**Figure S4C**), a mixture of maltose & lactose (**Figure S4D**) and actual perfusate sample as reference (**Figure S4E**). None of these were matching, indicating that the sample species remains to be confirmed. MS fragmentation matching scores suggested cellobiose, also known as GLCB1-4GLCB or cellose, in kidney perfusates under UR. Cellobiose, also classified as a reducing sugar, has been used as an indicator carbohydrate for Crohn’s disease and malabsorption syndrome ^29^. The origin of these disaccharides maybe most likely from external nutrients. For instance, 3-α-mannobiose, a member of the family of D-mannoses, is commonly found as a component of mannan, hemicellulose, and cellulose in dietary fibre ^30^. The structure of D-mannose is extremely similar to that of D-glucose and D-fructose, rendering the analytical determination considerably challenging. In general, the fact that disaccharides appear to accumulate in perfusates of UR NMP kidneys may reveal beneficial for controlling osmotic pressure ^31–33^, resulting in a more functional filtering and resorption function that maybe better maintained under UR NMP conditions.

Notably, we observed accumulated levels of N-(2-Acetamido)iminodiacetic acid (ADA) in NMP UR. ADA is a zwitterionic buffer often used to stabilise pH, but also a group P2 transcription activator, which enhances the expression of insulin from pancreatic cells and used for the treatment of diabetes through an epigenetic mechanism ^34,35^. ADA improves glucose metabolism and increases the glomerular filtration rate in diabetic patients. It is possible that some of the donor patients, whose discarded kidneys were used in this study, may have been subjected to treatment for diabetes (this patient information was not obtainable in the context of this study), inadvertently having beneficial effects on kidney function.

We also observed accumulation of purine derivatives as well as urea and glutamic acid in kidney UR-NMP (**Figure 4**). Urea recycling in kidneys has been described, leading to higher rates of potassium secretion ^36,37^. Also, urea-to-plasma concentration ratio affects eGFR ^38^. Urea levels may also affect urea transporter (UT-A1, UT-A3) feedback loops involved in regulating osmolarity, water reabsorption, vascular tension and enhanced kidney filtration function ^39,40^. Of note is a marked reduction in lactate (**Figure 4, 5E**) and more balanced glucose levels (**Figure 4, 5F**) in perfusates under UR as compared to NUR NMP conditions, further indicating a more balanced kidney energy homeostasis affecting both, oxidative phosphorylation and glycolysis pathways (**Figure 6**) ^41^. The exact underlying molecular details remain to be determined, and it may be that urine metabolic determinants other than urea itself, such as uric acid, creatinine and inorganic salts, may play a role. Interestingly, urine recirculation mediated changes in metabolic profiles show minimal evidence of uraemia, reflected by a slight increase in indol-3-acetic acid in UR NMP (**Figure 4A**), but may indicate a rewiring of the urea cycle that could ultimately feed back into carbohydrate and purine metabolic pathways (**Figure 6**) ^42^. The Urea and TCA cycles are closely interlinked as one of its nitrogen atoms is provided through transamination of oxalacetate to form aspartate, and, in turn, the delivery of fumarate to the TCA cycle (**Figure 6**) ^43^. Also, transamination of glutamate to 2-oxoglutarate may fuel the amino acid pool, consistent with an increase in L-leucine, L-methionine and N-acetyltyrosine levels (**Figure 4A**). The present study has limitations in that the sample size is small with the inclusion of eight discarded kidneys that were split between two groups (UR versus NUR NMP). Many clinical parameters, such as age, gender, DCD, DBD, CIT may contribute to additional variations within the perfusate metabolome, which is distinct from *in vivo* conditions ^44^. Clearly, independent validation and more detailed mechanistic and dynamic insights of these metabolic networks driven by UR may await specific metabolic flux studies performed under NMP conditions. Alternatively, urine fractionation could facilitate the identification of which molecular component(s) may contribute to increased NMP performance. Also, comparing the perfusate metabolic properties of kidney organs derived from living versus deceased donors may also reveal relevant insights.

In summary, our results describe differential perfusate metabolic profiles between UR and NUR NMP that may contribute to an elevated physiological state of kidney organs perfused with urine recirculation under normothermic conditions. This has implications, not only to help our molecular understanding, but also to justify the clinical implementation of UR to NMP protocols that lead to improved kidney functional preservation characteristics.

## Author contributions

A.W. conceptualised the study, collected discarded kidney samples and performed NMP to retrieve perfusate material. Z.Y., H.H., B.Y. and B.M.K. performed and analysed metabolomics data. A.W. and B.M.K. wrote the manuscript. All co-authors (A.W., Z.Y., H.H., L.F.M.L., B.Y., J.P.H, R.J.P., C.C.C., P.J.F. and B.M.K.) edited and provided feedback to the manuscript.

## Acknowledgements

We thank Dr Maria-Konstantina Ioannidi and Prof Maria Klapa (Patras University, Greece) for insightful discussions. This study was supported by the MRC/UKRI programme award QUOD-ATLAS (MR/R014132/1) to R.J.P. B.M.K. was supported by the Chinese Academy of Medical Sciences (CAMS) Innovation Fund for Medical Science (CIFMS), China (grant number: 2024-I2M-2-001-1).

## Data availability

All data sets including the metabolomics data will be made available upon request.

## Declaration of generative AI and AI-assisted technologies in the manuscript preparation process

During the preparation of this work, the authors have not used any AI nor any AI-assisted technologies.

## Abbreviations

ADA: N-(2-Acetamido)iminodiacetic acid
CIT: Cold ischemia time
DCD: Donation after circulatory death
DGF: Delayed graft function
ECD: Extended criteria death
eGFR: Estimated glomerular filtration rate
EI: Electron impact
ESRD: End stage renal disease
FDR: False discovery rate
GCxGC-qMS: Two-dimensional gas chromatography single quadrupole mass spectrometry
HMP: Hypothermic machine preservation
IQR: Interquartile range
IRAS: Integrated Research Application System
kPa: Kilo pascal
LoD: limit of detection
MSTFA: N-Methyl-N-trimethylsilyltrifluoroacetamide
MTBE: *Tert*-butyl methyl ether
NGAL: Neutrophil Gelatinase-Associated Lipocalin
NIST: National Institute of Standards and Technology
NMP: Normothermic perfusion
NUR: Urine replacement
REC: Research Ethics Service and Research Ethics Committees RRT Renal replacement therapy
SCS: Static cold storage
TCA: Citric acid cycle
TMCS: chlorotrimethylsilane
UR: Urine recirculation

## References

1. GBD Chronic Kidney Disease Collaboration. Global, regional, and national burden of chronic kidney disease, 1990-2017: a systematic analysis for the Global Burden of Disease Study 2017. Lancet. 2020;395(10225):709–733. doi:10.1016/S0140-6736(20)30045-3

2. Demiselle J, Augusto JF, Videcoq M, et al. Transplantation of kidneys from uncontrolled donation after circulatory determination of death: comparison with brain death donors with or without extended criteria and impact of normothermic regional perfusion. Transpl Int. 2016;29(4):432–442. doi:10.1111/tri.12722

3. Moers C, Smits JM, Maathuis MHJ, et al. Machine perfusion or cold storage in deceased-donor kidney transplantation. N Engl J Med. 2009;360(1):7–19. doi:10.1056/NEJMoa0802289

4. Treckmann J, Moers C, Smits JM, et al. Machine perfusion versus cold storage for preservation of kidneys from expanded criteria donors after brain death. Transpl Int. 2011;24(6):548–554. doi:10.1111/j.1432-2277.2011.01232.x

5. Jochmans I, Moers C, Smits JM, et al. Machine perfusion versus cold storage for the preservation of kidneys donated after cardiac death: a multicenter, randomized, controlled trial. Ann Surg. 2010;252(5):756–764. doi:10.1097/SLA.0b013e3181ffc256

6. Nicholson ML, Hosgood SA. Renal transplantation after ex vivo normothermic perfusion: the first clinical study. Am J Transplant. 2013;13(5):1246–1252. doi:10.1111/ajt.12179

7. Hosgood SA, Nicholson ML. The evolution of donation after circulatory death donor kidney repair in the United Kingdom. Curr Opin Organ Transplant. 2018;23(1):130–135. doi:10.1097/MOT.0000000000000477

8. Hosgood SA, Saeb-Parsy K, Wilson C, Callaghan C, Collett D, Nicholson ML. Protocol of a randomised controlled, open-label trial of ex vivo normothermic perfusion versus static cold storage in donation after circulatory death renal transplantation. BMJ Open. 2017;7(1):e012237. doi:10.1136/bmjopen-2016-012237

9. Hosgood SA, Nicholson ML. The first clinical case of intermediate ex vivo normothermic perfusion in renal transplantation. Am J Transplant. 2014;14(7):1690–1692. doi:10.1111/ajt.12766

10. DiRito JR, Hosgood SA, Tietjen GT, Nicholson ML. The future of marginal kidney repair in the context of normothermic machine perfusion. Am J Transplant. Published online June 7, 2018. doi:10.1111/ajt.14963

11. Klein Nulend R, Hameed A, Singla A, et al. Normothermic Machine Perfusion and Normothermic Regional Perfusion of DCD Kidneys Before Transplantation: A Systematic Review. Transplantation. Published online July 18, 2024. doi:10.1097/TP.0000000000005132

12. Weissenbacher A, Lo Faro L, Boubriak O, et al. Twenty-four-hour normothermic perfusion of discarded human kidneys with urine recirculation. Am J Transplant. Published online May 14, 2018. doi:10.1111/ajt.14932

13. Weissenbacher A, Huang H, Surik T, et al. Urine recirculation prolongs normothermic kidney perfusion via more optimal metabolic homeostasis-a proteomics study. Am J Transplant. Published online October 5, 2020. doi:10.1111/ajt.16334

14. Bon D, Billault C, Claire B, et al. Analysis of perfusates during hypothermic machine perfusion by NMR spectroscopy: a potential tool for predicting kidney graft outcome. Transplantation. 2014;97(8):810–816. doi:10.1097/TP.0000000000000046

15. Faucher Q, Alarcan H, Sauvage FL, et al. Perfusate Metabolomics Content and Expression of Tubular Transporters During Human Kidney Graft Preservation by Hypothermic Machine Perfusion. Transplantation. Published online April 20, 2022. doi:10.1097/TP.0000000000004129

16. Wijermars LGM, Schaapherder AF, de Vries DK, et al. Defective postreperfusion metabolic recovery directly associates with incident delayed graft function. Kidney Int. 2016;90(1):181–191. doi:10.1016/j.kint.2016.02.034

17. Guy AJ, Nath J, Cobbold M, et al. Metabolomic analysis of perfusate during hypothermic machine perfusion of human cadaveric kidneys. Transplantation. 2015;99(4):754–759. doi:10.1097/TP.0000000000000398

18. Nath J, Guy A, Smith TB, et al. Metabolomic perfusate analysis during kidney machine perfusion: the pig provides an appropriate model for human studies. PLoS ONE. 2014;9(12):e114818. doi:10.1371/journal.pone.0114818

19. Weissenbacher A. Normothermic Kidney Preservation. http://purl.org/dc/dcmitype/Text. University of Oxford; 2018. Accessed June 7, 2020. https://ora.ox.ac.uk/objects/uuid:57ae08d0-bf5c-422d-af85-893e15e6ec7c

20. Yu Z, Huang H, Reim A, et al. Optimizing 2D gas chromatography mass spectrometry for robust tissue, serum and urine metabolite profiling. Talanta. 2017;165:685–691. doi:10.1016/j.talanta.2017.01.003

21. Huang H, van Dullemen LFA, Akhtar MZ, et al. Proteo-metabolomics reveals compensation between ischemic and non-injured contralateral kidneys after reperfusion. Sci Rep. 2018;8(1):8539. doi:10.1038/s41598-018-26804-8

22. Beens J, Adahchour M, Vreuls RJ, van Altena K, Brinkman UA. Simple, non-moving modulation interface for comprehensive two-dimensional gas chromatography. J Chromatogr A. 2001;919(1):127–132.

23. Tyanova S, Temu T, Sinitcyn P, et al. The Perseus computational platform for comprehensive analysis of (prote)omics data. Nat Methods. 2016;13(9):731–740. doi:10.1038/nmeth.3901

24. Akhtar MZ, Huang H, Kaisar M, et al. Using an Integrated -Omics Approach to Identify Key Cellular Processes That Are Disturbed in the Kidney After Brain Death. American Journal of Transplantation. 2016;16(5):1421–1440. doi:10.1111/ajt.13626

25. Vaughan RH, Kresse JC, Farmer LK, et al. Cytoskeletal protein degradation in brain death donor kidneys associates with adverse post-transplant outcomes. Am J Transplant. Published online December 8, 2021. doi:10.1111/ajt.16912

26. L. L. van Leeuwen, L. H. Venema, H. G.D. Leuvenink, B. M. Kessler. Doxycycline alters renal secretome and tissue degradome during hypothermic machine perfusion leading to delayed onset of fibrosis. 2021. Under review.

27. Udzik J, Pacholewicz J, Biskupski A, et al. Higher perfusion pressure and pump flow during cardiopulmonary bypass are beneficial for kidney function–a single-centre prospective study. Front Physiol. 2024;15. doi:10.3389/fphys.2024.1257631

28. Wei J, Song J, Jiang S, et al. Role of intratubular pressure during the ischemic phase in acute kidney injury. Am J Physiol Renal Physiol. 2017;312(6):F1158–F1165. doi:10.1152/ajprenal.00527.2016

29. Welcker K, Martin A, Kölle P, Siebeck M, Gross M. Increased intestinal permeability in patients with inflammatory bowel disease. Eur J Med Res. 2004;9(10):456–460.

30. Hu X, Shi Y, Zhang P, Miao M, Zhang T, Jiang B. d-Mannose: Properties, Production, and Applications: An Overview. Comprehensive Reviews in Food Science and Food Safety. 2016;15(4):773–785. doi:10.1111/1541-4337.12211

31. Wen L, Li Y, Li S, Hu X, Wei Q, Dong Z. Glucose Metabolism in Acute Kidney Injury and Kidney Repair. Front Med (Lausanne). 2021;8:744122. doi:10.3389/fmed.2021.744122

32. Dickenmann M, Oettl T, Mihatsch MJ. Osmotic nephrosis: acute kidney injury with accumulation of proximal tubular lysosomes due to administration of exogenous solutes. Am J Kidney Dis. 2008;51(3):491–503. doi:10.1053/j.ajkd.2007.10.044

33. Pessoa TD, Campos LCG, Carraro-Lacroix L, Girardi ACC, Malnic G. Functional role of glucose metabolism, osmotic stress, and sodium-glucose cotransporter isoform-mediated transport on Na+/H+ exchanger isoform 3 activity in the renal proximal tubule. J Am Soc Nephrol. 2014;25(9):2028–2039. doi:10.1681/ASN.2013060588

34. Singh R, Chandel S, Dey D, et al. Epigenetic modification and therapeutic targets of diabetes mellitus. Biosci Rep. 2020;40(9):BSR20202160. doi:10.1042/BSR20202160

35. Mahne CW, Salah-Uddin H. Treatment of diabetes and associated metabolic conditions with epigenetic modulators. Published online August 16, 2018. Accessed September 29, 2024. https://patents.google.com/patent/WO2018148206A1/en/

36. Halperin ML, Gowrishankar M, Mallie JP, Sonnenberg H, Oh M. Urea recycling: an aid to the excretion of potassium during antidiuresis. Nephron. 1996;72(4):507–511. doi:10.1159/000188930

37. Kamel KS, Halperin ML. Intrarenal urea recycling leads to a higher rate of renal excretion of potassium: an hypothesis with clinical implications. Curr Opin Nephrol Hypertens. 2011;20(5):547–554. doi:10.1097/MNH.0b013e328349b8f9

38. Petrovic D, Bankir L, Ponte B, et al. The urine-to-plasma urea concentration ratio is associated with eGFR and eGFR decline over time in a population cohort. Nephrol Dial Transplant. 2023;39(1):122–132. doi:10.1093/ndt/gfad131

39. Esteva-Font C, Anderson MO, Verkman AS. Urea transporter proteins as targets for small-molecule diuretics. Nat Rev Nephrol. 2015;11(2):113–123. doi:10.1038/nrneph.2014.219

40. Yu L, Liu T, Fu S, et al. Physiological functions of urea transporter B. Pflugers Arch - Eur J Physiol. 2019;471(11-12):1359–1368. doi:10.1007/s00424-019-02323-x

41. Arykbaeva AS, De Vries DK, Doppenberg JB, et al. Metabolic needs of the kidney graft undergoing normothermic machine perfusion. Kidney International. 2021;100(2):301–310. doi:10.1016/j.kint.2021.04.001

42. Wang Z, Gu Z, Shen Y, et al. The Natural Product Resveratrol Inhibits Yeast Cell Separation by Extensively Modulating the Transcriptional Landscape and Reprogramming the Intracellular Metabolome. PLoS One. 2016;11(3):e0150156. doi:10.1371/journal.pone.0150156

43. Yoshimi N, Futamura T, Kakumoto K, et al. Blood metabolomics analysis identifies abnormalities in the citric acid cycle, urea cycle, and amino acid metabolism in bipolar disorder. BBA Clin. 2016;5:151–158. doi:10.1016/j.bbacli.2016.03.008

44. Ogurlu B, van Furth LA, Zuo Y, et al. Kidney Metabolism During Normothermic Machine Perfusion Differs Substantially From In Vivo Conditions. Transplantation. 2025 Dec 3. doi: 10.1097/TP.0000000000005586.

