## SUPPLEMENTAL FILES for "Urine Recirculation During Normothermic Kidney Preservation Improves Energy Balance Involving the Urea and TCA Cycles"

### 1. SUPPLEMENTAL FIGURES

**Figure S1: Global metabolic profiles of NMP NUR versus UR kidney perfusate.** (**A**) Correlation analysis. (**B**) Box plot analysis to assess sample normalisation.

**Figure S2: Metabolic profiles in kidney NMP perfusates distinguish mono- and di-saccharides.** (**A**) 2-dimensional gas chromatography (GCxGC) plot displaying glucose (monosaccharide) and disaccharide metabolic species. (**B**) 3-dimensional representation of the GCxGC chromatogram displayed above.

**Figure S3: MS fragment spectra discriminate glucose from disaccharides.** (**A**) Electron impact (EI) MS spectrum for glucose and derivatives. Representative data derived from sample kidney 35 24hr NMP UR (140618AM26NTX3524h1ul20split003). (**B**) EI MS spectra for disaccharides. Representative data derived from sample kidney 35 24hr NMP UR (140618AM26NTX35-24h1ul20split003). Both EI MS spectra were generated using the GCMSSolution software (see Materials and Methods).

**Figure S4: Disaccharide identification using standards.** (**A**) Comparison of GCxGC chromatograms between kidney NMP UR perfusate (purple) and glucose standard (black). (**B**) Comparison of GCxGC chromatograms between kidney NMP

UR perfusate (purple) and lactose standard (black). (**C**) Comparison of GCxGC chromatograms between kidney NMP UR perfusate (purple) and maltose standard (black). (**D**) Comparison of GCxGC chromatograms between kidney NMP UR perfusate (purple) and a lactose and maltose standard mix (black). (**E**) GCxGC chromatogram of kidney NMP UR perfusate alone as reference.

### 2. SUPPLEMENTAL TABLES

**Table S1: Metabolites correlating with perfusion conditions. (A)** Correlation of donor and perfusion characteristics with metabolites **(B)** Differences in the perfusate metabolome of *UR and *NUR kidneys after 1 hour of NMP. **(C)** Differences in the perfusate metabolome of *UR and *NUR kidneys after 6 hours of NMP**.**

**Figure S1**

**A**

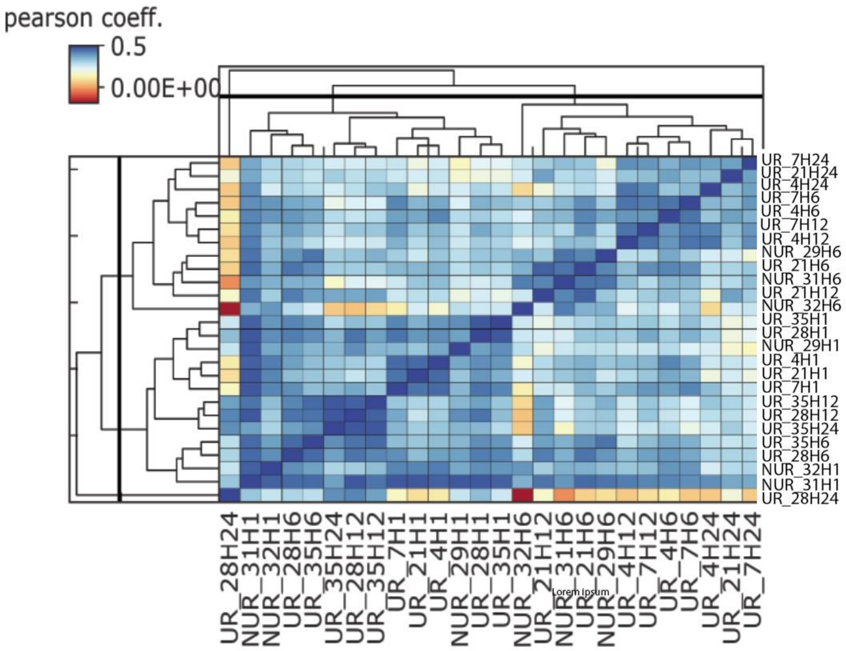
pearson coeff.
0.5
0.00E+00
UR_7H24
UR_21H24
UR_4H24
UR_7H6
UR_4H6
UR_7H12
UR_4H12
NUR_29H6
UR_21H6
NUR_31H6
UR_21H12
NUR_32H6
UR_35H1
UR_28H1
NUR_29H1
UR_4H1
UR_21H1
UR_7H1
UR_35H12
UR_28H12
UR_35H24
UR_35H6
UR_28H6
NUR_32H1
NUR_31H1
UR_28H24

**B**

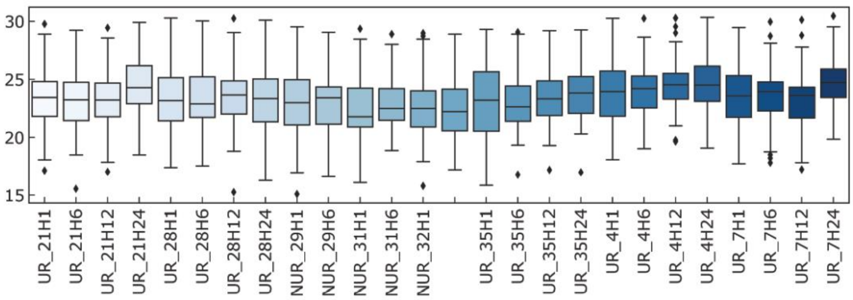
30
25
20
15
UR_21H1
UR_21H6
UR_21H12
UR_21H24
UR_28H1
UR_28H6
UR_28H12
UR_28H24
NUR_29H1
NUR_29H6
NUR_31H1
NUR_31H6
NUR_32H1
UR_35H1
UR_35H6
UR_35H12
UR_35H24
UR_4H1
UR_4H6
UR_4H12
UR_4H24
UR_7H1
UR_7H6
UR_7H12
UR_7H24

**Figure S2**

A

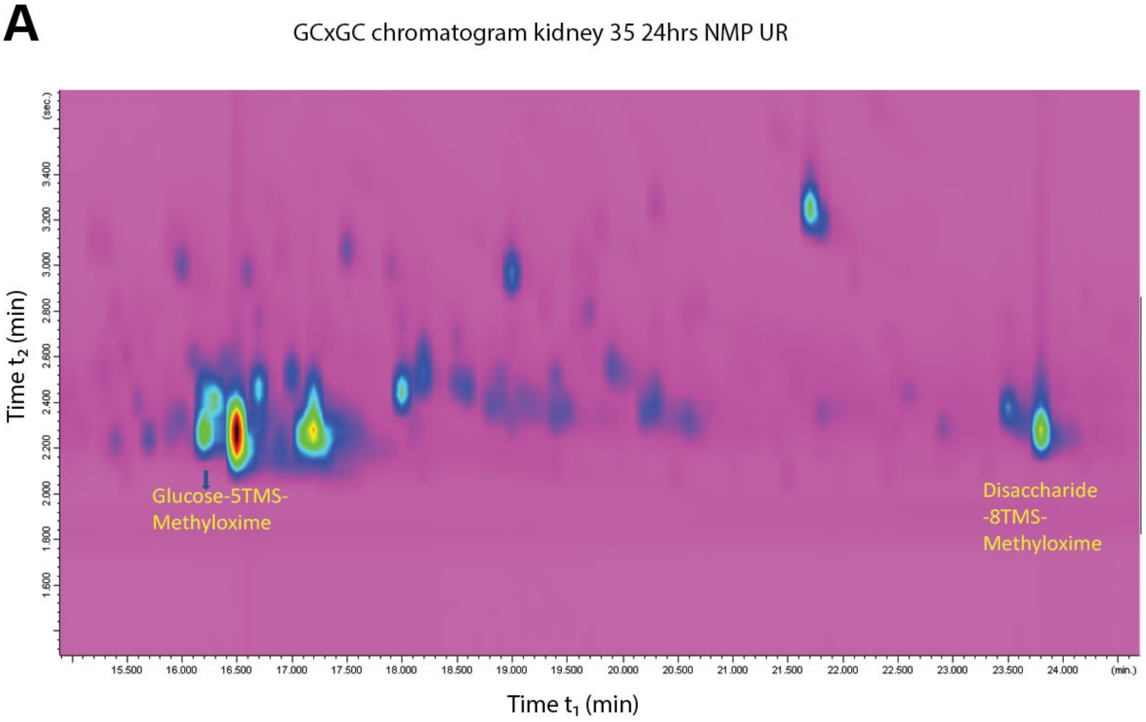
GCxGC chromatogram kidney 35 24hrs NMP UR
Time $t_2$ (min)
(sec.)
3.400
3.200
3.000
2.800
2.600
2.400
2.200
2.000
1.800
1.600
Glucose-5TMS-
Methyloxime
Disaccharide
-8TMS-
Methyloxime
15.500
16.000
16.500
17.000
17.500
18.000
18.500
19.000
19.500
20.000
20.500
21.000
21.500
22.000
22.500
23.000
23.500
24.000
(min.)
Time $t_1$ (min)

B

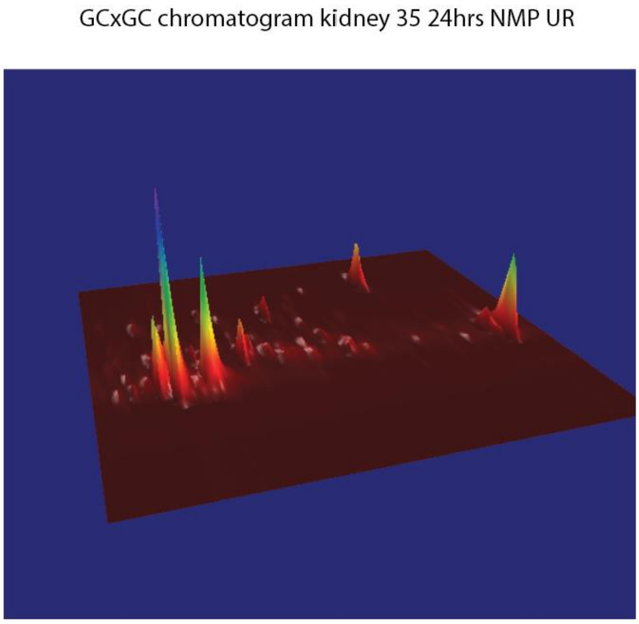
GCxGC chromatogram kidney 35 24hrs NMP UR

**Figure S3**

**A** Similarity search for Glucose from GCMSSolution

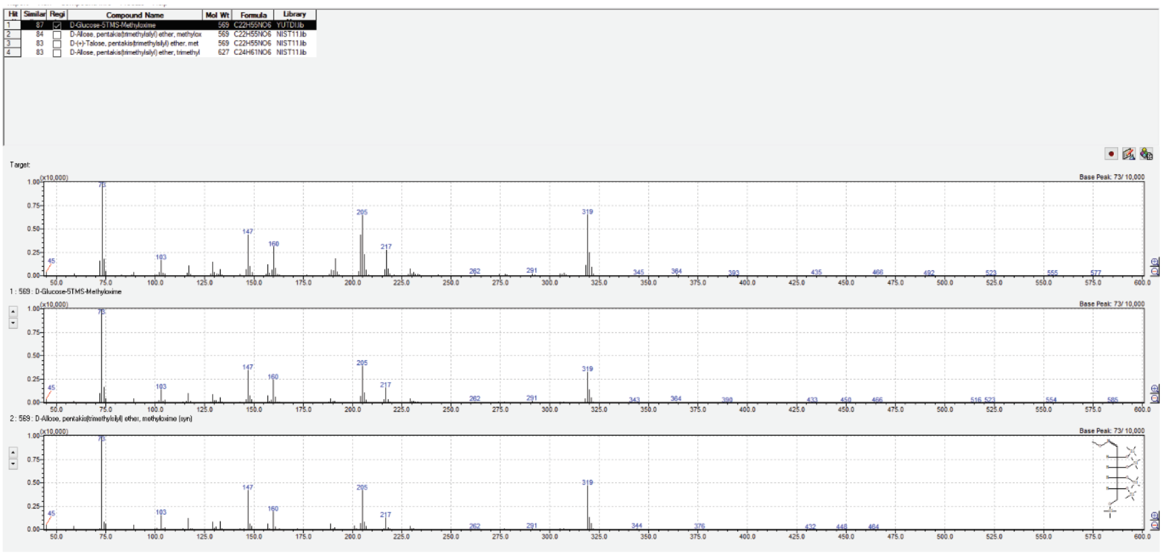


**B** Similarity Search for Disaccharide from GCMSSolution

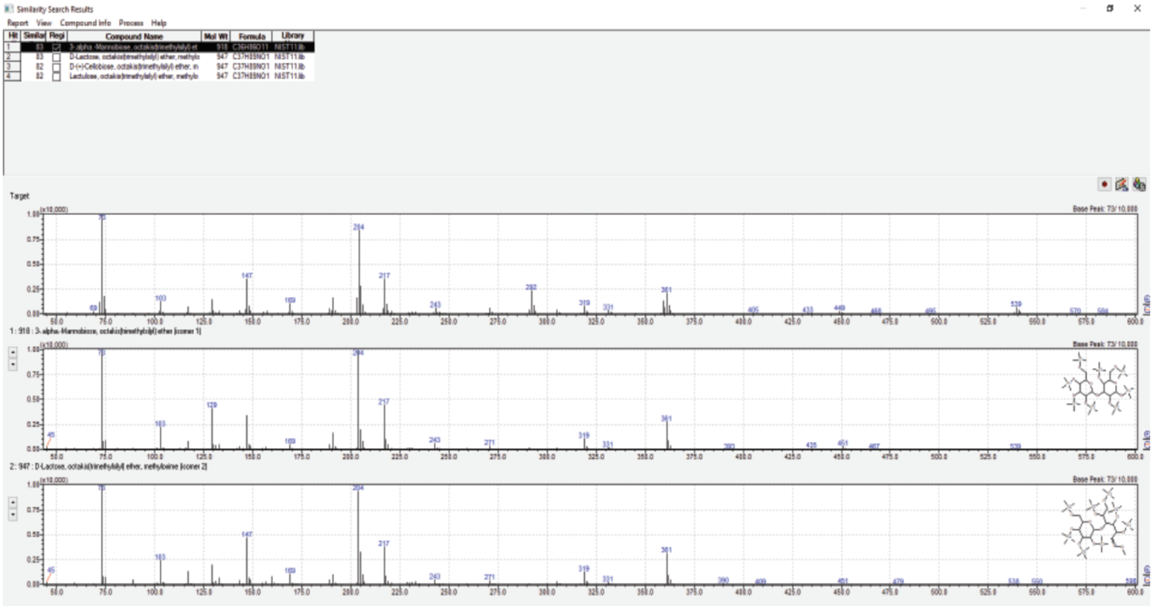

**Figure S4**

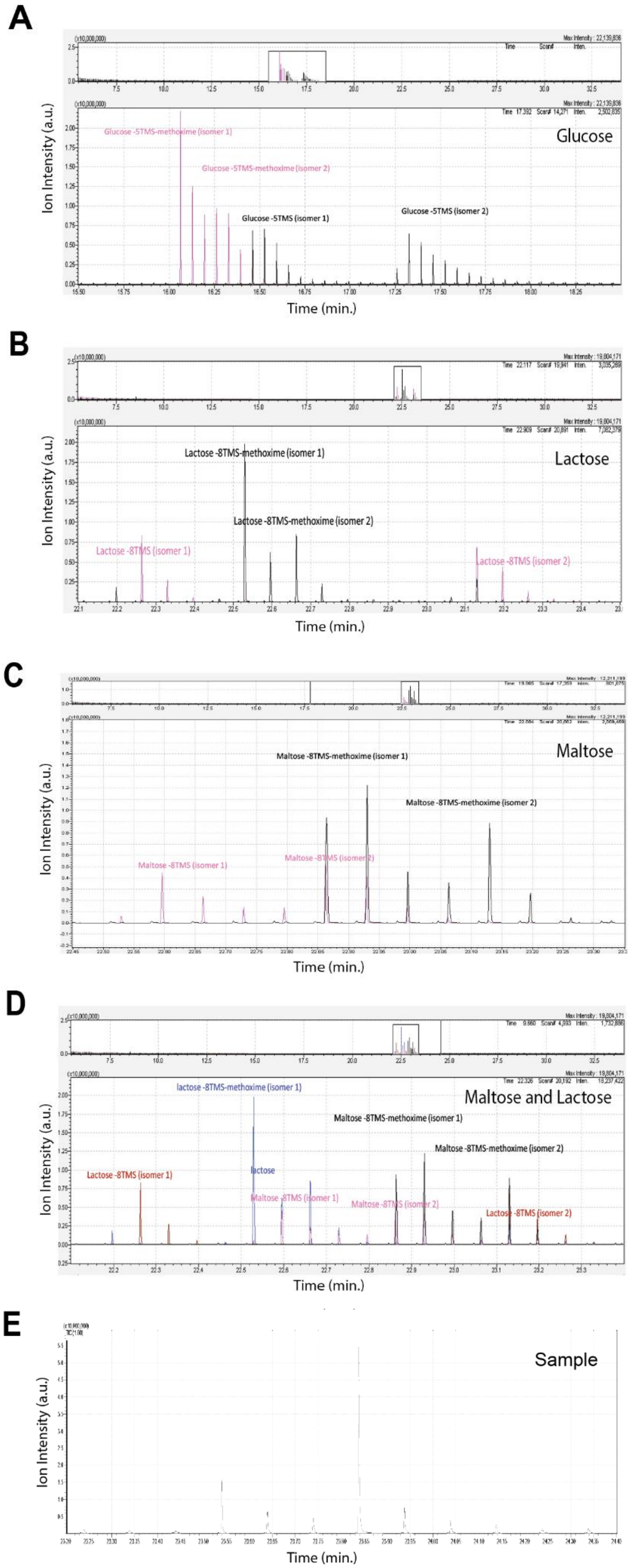
A
Ion Intensity (a.u.)
Glucose
Glucose -5TMS-methoxime (isomer 1)
Glucose -5TMS-methoxime (isomer 2)
Glucose -5TMS (isomer 1)
Glucose -5TMS (isomer 2)
Time (min.)
B
Ion Intensity (a.u.)
Lactose
Lactose -8TMS-methoxime (isomer 1)
Lactose -8TMS-methoxime (isomer 2)
Lactose -8TMS (isomer 1)
Lactose -8TMS (isomer 2)
Time (min.)
C
Ion Intensity (a.u.)
Maltose
Maltose -8TMS-methoxime (isomer 1)
Maltose -8TMS-methoxime (isomer 2)
Maltose -8TMS (isomer 1)
Maltose -8TMS (isomer 2)
Time (min.)
D
Ion Intensity (a.u.)
Maltose and Lactose
lactose -8TMS-methoxime (isomer 1)
Maltose -8TMS-methoxime (isomer 1)
Maltose -8TMS-methoxime (isomer 2)
Lactose -8TMS (isomer 1)
lactose
Maltose -8TMS (isomer 1)
Maltose -8TMS (isomer 2)
Lactose -8TMS (isomer 2)
Time (min.)
E
Ion Intensity (a.u.)
Sample
Time (min.)

**Table S1A: Correlation of donor and perfusion characteristics with metabolites**

| Metabolite | p value | Spearman's rho | Donor characteristics |
|---|---|---|---|
| Lactate | 0.02 | 0.81 | eGFR |
| Gluconate | 0.03 | 0.79 | eGFR |
| L-tyrosine | 0.01 | 0.84 | Duration of NMP |
| Phosphoglycerol | 0.01 | 0.85 | Duration of NMP |
| L-tyrosine | 0.007 | -0.88 | Urine output during NMP |
| Phosphoglycerol | 0.01 | -0.86 | Urine output during NMP |
| Disaccharide | 0.04 | 0.87 | UR NMP |
| Gluconate | 0.04 | 0.85 | UR NMP |
| Phosphoglycerol | 0.04 | 0.85 | UR NMP |
| Lactate | 0.02 | -0.83 | Median arterial flow during NMP |
| Lactate | 0.05 | 0.72 | Median (IQR) $pCO_2$ levels during NMP |

**Table S1B: Differences in the perfusate metabolome of *UR and *NUR kidneys after 1 hour of NMP**

| Metabolite | p value | Mean rank of UR hour 1 | Mean rank of NUR hour 1 | Mean rank diff. | Mann-Whitney U | q value |
|---|---|---|---|---|---|---|
| .β.-D-Allopyranose | <0,000001 | 22.5 | 6.5 | 16 | 0 | <0,000001 |
| D-glucose | | | | | | |
| 2,6-dihydroxy-7H-purine | <0,000001 | 22.5 | 6.5 | 16 | 0 | <0,000001 |
| 6-hydroxy-9H-purine | <0,000001 | 22.5 | 6.5 | 16 | 0 | <0,000001 |
| L-alanine | <0,000001 | 22.5 | 6.5 | 16 | 0 | <0,000001 |
| D-(+)-Cellobiose (disaccharide) | <0,000001 | 22.5 | 6.5 | 16 | 0 | <0,000001 |
| Lactate | <0,000001 | 22.5 | 6.5 | 16 | 0 | <0,000001 |
| 3-hydroxybutyrate | <0,0003 | 22.5 | 6.5 | 16 | 0 | <0,000001 |
| Urea | <0,000001 | 22.5 | 6.6 | 16 | 1 | <0,000001 |

*UR= urine recirculation; NUR= no urine recirculation/urine replacement with Ringer's lactate

**Table S1C: Differences in the perfusate metabolome of *UR and *NUR kidneys after 6 hours of NMP**

| Metabolite | p value | Mean rank of UR hour 6 | Mean rank of NUR hour 6 | Mean rank diff. | Mann-Whitney U | q value |
|---|---|---|---|---|---|---|
| .β.-D-galactofuranose | <0,000001 | 22.5 | 6.5 | 16 | 0 | <0,000001 |
| Disaccharide (D-(+)-Cellobiose) | <0,000001 | 22.5 | 6.5 | 16 | 0 | <0,000001 |
| D-gluconate | <0,001 | 22.5 | 6.5 | 16 | 0 | <0,000001 |
| Glutamate | <0,000001 | 22.5 | 6.5 | 16 | 0 | <0,000001 |
| Lactate | <0,000001 | 22.5 | 6.5 | 16 | 0 | <0,000001 |
| L-serine | <0,001 | 22.5 | 6.5 | 16 | 0 | <0,000001 |
| Glycine | <0,004 | 22.5 | 6.5 | 16 | 0 | <0,000001 |
| L-aspartate | <0,0001 | 22.5 | 6.5 | 16 | 0 | <0,000001 |
| Urea | <0,000001 | 22.5 | 6.5 | 16 | 0 | <0,000001 |

*UR= urine recirculation; NUR= no urine recirculation/urine replacement with Ringer's lactate
